# *Junb* acts downstream of the *Il-11/Stat3* signaling axis to limit tissue damage induced fibro-inflammation during regeneration

**DOI:** 10.64898/2026.08.04.742722

**Authors:** Julia Ishaque, Eric Bienert, Amrutha Manikandan, Srinivas Allanki, Laia Cañes-Esteve, Jochen Pöling, Stefan Günther, Didier Y.R. Stainier, Samuel T. Sossalla, Arica Beisaw, Sven Reischauer

## Abstract

Inflammation is essential for regeneration yet can also drive fibroinflammatory remodeling; what determines these opposing outcomes remains unclear. Comparative transcriptomic analyses revealed that injured mouse hearts activated a broad inflammatory program, whereas zebrafish hearts mounted a restricted response characterized by selective *il11* induction. This divergence extended across tissues and species: the non-regenerative mammalian injuries examined shared an inflammatory signature distinct from regenerative vertebrate contexts. We identified the AP-1 transcription factor Junb as an Il-11–Stat3-dependent regulator that restrains inflammation during fin fold regeneration. Combined loss of *junba* and *junbb* amplified a mammalian-like inflammatory program, increased neutrophil recruitment and fibroinflammatory gene expression, and reduced proliferation and regenerative outgrowth. Strikingly, dexamethasone or ibuprofen substantially restored regeneration in Junb deficient zebrafish larvae, demonstrating that hyperinflammation is a major determinant of regenerative failure. Thus, the Il-11–Stat3–Junb axis maintains a regeneration permissive inflammatory state preventing a regenerative response from shifting toward mammalian-like fibroinflammation.

## INTRODUCTION

The capacity for tissue regeneration exhibits remarkable variation among vertebrate species, ranging from limited capabilities in mammals, including humans, to exceptional regenerative abilities in some aquatic vertebrates including bony fish and amphibians (e.g., zebrafish and axolotl). While mammals often respond to tissue damage with fibrotic scarring, resulting in impaired organ function, regenerative species can fully restore tissues, including appendages and the heart without permanent scar formation. This dichotomous nature suggests a causal relationship between scar formation and reduced regenerative potential. Hence, understanding the underlying cellular and molecular mechanisms has been a subject of intense scientific investigation (1).

A variety of factors, pathways, and cellular processes have been identified as important for successful regeneration. While dedifferentiation, proliferation, and cell migration are key cellular processes in the formation and patterning of the regenerate, activation of the local immune system and the recruitment of immune cells represent additional critical components of the regenerative response in the heart and other tissues (2). Notably, the ablation of immune cells including macrophages, granulocytes, and neutrophils attenuates or even completely blocks regeneration (3). However, tissue inflammation is known to exert detrimental effects on regeneration. Prolonged or excessive inflammation can limit the regenerative potential and induce fibrotic remodeling, a process termed fibro-inflammation (4,5). Taken together, these findings suggest that inflammation represents a default response to tissue injury; however, its qualitative and quantitative characteristics may differ substantially between regenerative and non-regenerative species. Moreover, successful regeneration clearly requires that this inflammatory response be tightly controlled, both in magnitude and duration. Despite this central role, mechanistic insight into the qualitative differences that distinguish a pro-regenerative from a non-regenerative injury-induced inflammatory response, as well as into the mechanisms that regulate this transition, remains conspicuously limited. AP-1 comprises of family of dimeric transcription factor complexes that are rapidly induced as immediate-early genes in response to diverse extracellular signals, and is mainly composed of members of the JUN, FOS, ATF and MAF gene families. AP-1 induction has been observed in border-zone cardiomyocytes after myocardial infarction in mice, as well as after cardiac injury in zebrafish. Notably, AP-1 transcription factors have been shown to play key roles in the response to tissue injury and in cardiomyocyte regeneration in zebrafish. Among AP-1 family members, Junb was specifically shown to increase chromatin accessibility in regenerating cardiomyocytes after zebrafish cardiac injury, suggesting that AP-1 activity may contribute not only to the transcriptional response to injury, but also to the establishment of a permissive chromatin landscape during regeneration (6).

The zebrafish heart and fins are established models for studying regenerative biology, offering insights into tissue specific but also universal mechanisms underlying tissue regeneration (7,8). Heart and fin regeneration in zebrafish exhibit very similar innate immune dynamics. This conserved early sequence indicates shared inflammatory requirements for regeneration across tissues, with macrophages contributing to inflammation resolution and the establishment of a pro-regenerative environment. In contrast to regenerating zebrafish hearts surprisingly little is known regarding the early phases in mouse hearts after tissue damage.

Here, we systematically compare early injury-induced inflammation in mouse and zebrafish hearts using comparative transcriptomics. We show that inflammatory gene expression, particularly within the Interleukin-6 family differs substantially with non-regenerative mice showing a rapid activation of a highly inflammatory gene program while regenerative zebrafish hearts do not. Using genetic loss-of-function models in combination with high resolution imaging expression analysis and pharmacological experiments, we show that, in zebrafish, Interleukin-11, which is rapidly upregulated after injury, acts upstream of a *junb* mediated signaling cascade to limit excess-inflammation and promote regeneration.

## RESULTS

### Early cardiac injury responses diverge between regenerative and fibrotic models

Adult zebrafish fully regenerate damaged myocardium after injury, whereas adult mice respond to myocardial infarction with fibrotic remodeling and the formation of a functionally inert scar (9,10). To investigate whether these divergent outcomes are already reflected during the acute phase after cardiac injury, we compared transcriptional responses in mouse and zebrafish hearts 24 hours after injury. To this end, we isolated RNA from injured mouse hearts after LAD ligation and from injured zebrafish ventricles after cardiac cryoinjury, followed by bulk RNA sequencing and comparative bioinformatic analysis (Fig. 1A).

**Figure 1:**
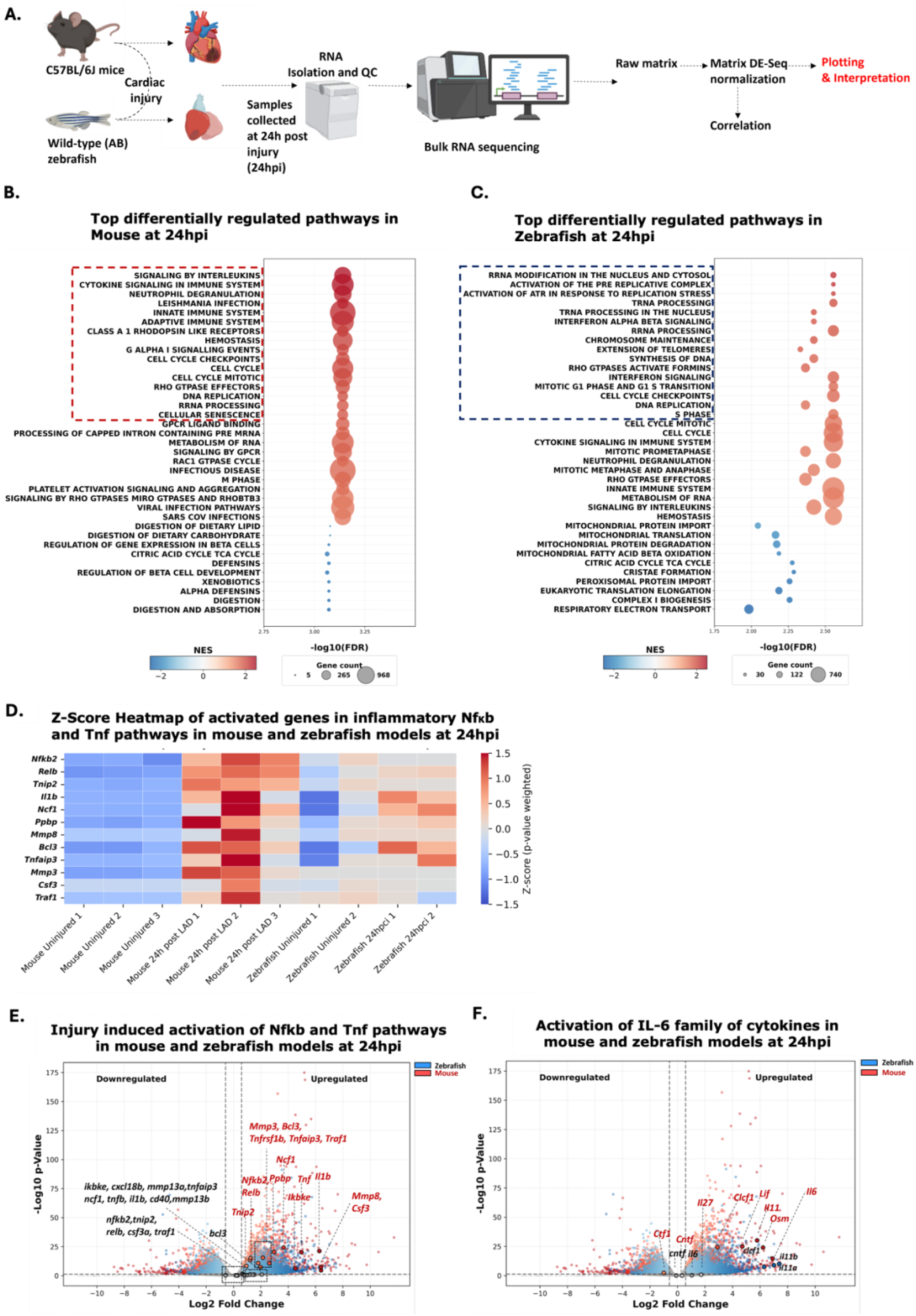
Transcriptional response to injury is differentially regulated in mice vs. zebrafish. **A.** Schematic depiction of the pipeline from injury (LAD ligation in mice vs. cryoinjury in zebrafish) to RNA sequencing. The LAD ligation was performed on three biological replicates of adult C57/Bl6J mice each for the control and injured conditions (n=3). The zebrafish cardiac cryoinjury was performed on 2 replicates of 5 pooled hearts, similarly 5 hearts were pooled for 2 replicates of the control conditions. Following injury, the samples were subjected to RNA isolation and sequencing. **B-C** Gene set enrichment analysis (GSEA) was performed on **B.** the mice dataset and **C.** the zebrafish dataset. The fgsea package from R and REACTOME was used for pathway analysis. The top 25 pathways that were differentially regulated with a p value and FDR < 0.02 were chosen in the representative image **D.** Transcriptional profile of differentially regulated genes involved in inflammation, in the *Nfkb* and *T*nf signalling pathways. Heatmap represent the z-score weighted for the p-value of each gene. **E.** Transcriptional profile of differentially regulated genes involved in inflammation, in the *Tnf* and *Nfkb* pathway. Volcano plot wwas generated using matplotlib with the threshold of |log2FC| = 0.585 and p-value < 0.05. Data for mice and zebrafish are represented in red and blue respectively, with gray coloration for unregulated genes. **F.** Transcriptional profile of differentially regulated genes in the *Il6* family of cytokines. Volcano plot wwas generated using matplotlib with the threshold of |log2FC| = 0.585 and p-value < 0.05. Data for mice and zebrafish are represented in red and blue respectively, with gray coloration for unregulated genes.

Gene set enrichment analysis revealed that both species activate a rapid injury response characterized by inflammatory and hemostatic programs (Fig. 1B, C). In both mouse and zebrafish, gene sets associated with cytokine and interleukin signaling, innate immune activation, neutrophil degranulation, interferon responses, and antiviral responses were significantly enriched, indicating that acute immune activation is a shared feature of cardiac injury. However, the zebrafish response was additionally characterized by enrichment of biosynthetic and proliferative pathways, including RNA metabolism, rRNA and tRNA processing, DNA replication, S phase, G2-M checkpoint, and ATR-mediated replication stress responses (Fig. 1C). In contrast, the mouse transcriptional profile was dominated by inflammatory signaling and stress-associated programs, including cellular senescence (Fig. 1B). Thus, regenerative and fibrotic cardiac injury responses share acute immune activation but diverge in the relative prominence of inflammatory versus proliferative transcriptional programs.

To further resolve this divergence, we examined genes associated with NF-κB and TNF signaling, two central inflammatory modules induced after tissue damage. At 24 hours after cardiac injury, mouse hearts showed a markedly stronger pro-inflammatory transcriptional response than zebrafish hearts (Fig. 1D, E). In mouse, canonical NF-κB/TNF-associated transcripts, including *Tnf*, *Il-1b*, *Tnfaip3*, *Tnfrsf1b*, *Traf1*, *Relb*, and *Nfκb2*, were robustly induced, together with inflammatory mediators and tissue-remodeling genes such as *Csf3*, *Mmp3*, and *Mmp8* (Fig. 1D, E). In contrast, zebrafish displayed an attenuated response within the same inflammatory module, with weaker induction and, for a subset of genes, reduced expression after injury (Fig. 1D, E). These data are consistent with previous descriptions of amplified inflammatory reactions in mammals after injury in diverse tissues (11) and indicate that early cardiac injury responses differ quantitatively and qualitatively between regenerative and fibrotic models (11). We next examined the regulation of IL-6 family cytokines as representative mediators of acute inflammatory signaling. This analysis revealed a restricted activation of IL-6 family ligands in zebrafish compared with mouse (Fig. 1F). In zebrafish, induction within the applied thresholds was largely restricted to *clcf1* and *il-11*, with *il-11* showing particularly robust upregulation, including both paralogues. In contrast, mouse hearts exhibited a broader IL-6 family response. In addition to *Il-11* and *Clcf1*, the ligands *Lif* and *Osm* were upregulated in mouse, and the canonical pro-inflammatory cytokine *Il6* showed the strongest induction (Fig. 1F).

Together, these data show that regenerative and nonregenerative cardiac injury responses are not distinguished by the presence or absence of inflammation, but by the magnitude and composition of the early inflammatory program. Extending this comparison to additional regenerative and non-regenerative injury contexts such as the adult zebrafish fin, larval zebrafish tail, and axolotl limb, as well as mouse digit-tip and porcine skin injury models further supported this pattern (suppl. Fig. 1). Regenerative systems exhibited a more restrained inflammatory architecture, whereas non-regenerative mammalian injury models showed stronger activation of canonical inflammatory nodes associated with *Il-1b*, *Il-6*, *Tnf*, *Tlr*, and *Nf-κB* signaling (suppl. Fig. 1). Thus, early injury-induced inflammatory signaling is selectively channeled in regenerative contexts, with *il-11* representing a prominent IL-6 family cytokine induced in the regenerating zebrafish heart.

### Il-11 signaling activates junba and junbb during the early injury response

Interleukin-11 (Il-11) has emerged as an important component of the injury response across species and tissues (12). While studies in mammalian systems have linked IL-11 to fibrotic and pro-inflammatory remodeling, other work has described tissue-protective or pro-regenerative functions (7,13). We previously provided genetic evidence that Il-11 signaling is required for scar-free regeneration across multiple tissues in zebrafish (7). These findings raised the possibility that, in regenerative contexts, Il-11 not only promotes tissue restoration but also restricts excessive inflammatory and fibrotic responses through downstream transcriptional regulators. Among the candidate mediators of such a response, we focused on Junb. *Junb* is rapidly induced in the injured zebrafish heart and has been implicated in regeneration-associated transcriptional programs in both fins and hearts in zebrafish (6,14). In addition, some studies in mammalian systems have linked JUNB to attenuation of IL-6 and TNF signaling, suggesting that Junb may act as a conserved modulator of inflammatory output (15,16). To test whether Junb functions downstream of Il-11 signaling after tissue damage, we used the zebrafish larval fin fold as a tractable model of scar-free regeneration. Although anatomically simple, the fin fold undergoes a stereotypical regenerative sequence involving wound closure, immune cell recruitment, blastema formation, and regenerative outgrowth, thereby recapitulating central features of injury-induced regeneration observed in more complex tissues (8). Consistently, cross-context analysis supported the use of larval zebrafish appendage regeneration as a mechanistic model to investigate regenerative inflammatory responses (suppl. Fig. 1), and *il11ra* mutant larvae failed to regenerate the fin fold, in line with the previously described requirement for Il-11 signaling during adult zebrafish tissue regeneration (7). Thus, this model provides a suitable system to examine how IL-11-dependent signaling regulates injury-induced responses during regeneration (7).

We first examined the expression of the two zebrafish *junb* paralogues, *junba* and *junbb*, after fin fold amputation using HCR-FISH. Both genes were rapidly induced after injury but displayed partially distinct expression domains: *junba* expression was enriched in epithelial layers, whereas *junbb* prominently marked the wound-adjacent central fin fold region, with partially overlapping expression at the wound site (Fig. 2A). To determine whether this induction depends on Il-11 signaling, we compared *junba* and *junbb* expression in wild-type and *il11ra* mutant larvae at early time points after amputation. In wild-type larvae, both paralogues were induced within 30 min after injury and remained detectable during the early regenerative response. In contrast, *il11ra^-/-^* mutants showed markedly reduced expression of both *junba* and *junbb* across the analyzed time points, including 0.5, 1, 3, and 6 hpa (Fig. 2A). These data indicate that Il-11 receptor signaling is required for the proper early activation of both *junb* paralogues after tissue damage.

**Figure 2:**
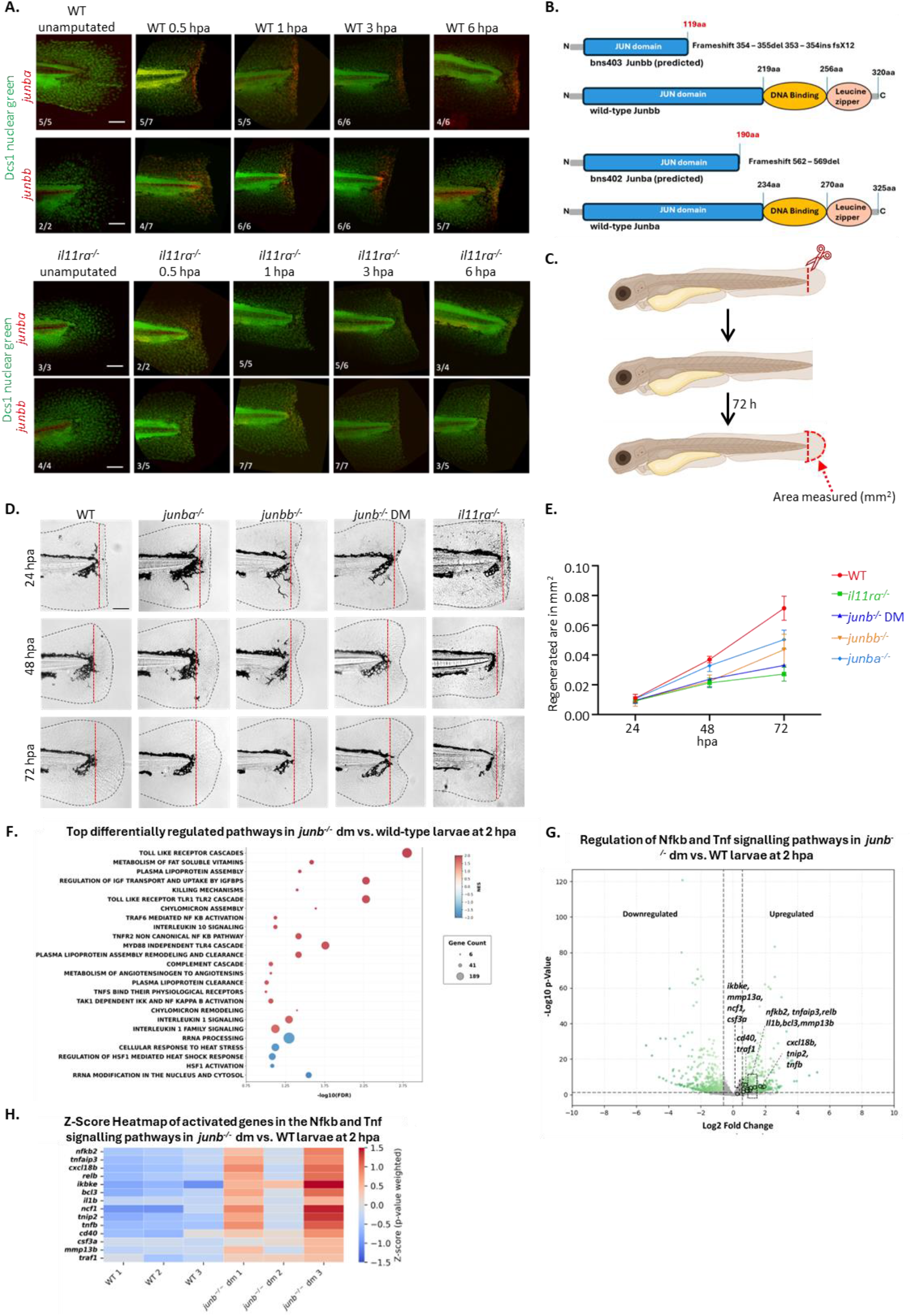
Expression of *junba* & *junbb* is triggered by injury, facilitated by Interleukin 11 signalling and cruical for tissue regeneration. **A.** Whole mount HCR-FISH for *junba* (red, upper rows) and *junbb* (red, lower rows) and nuclear counter stain (green) in WT (upper two rows) and *il11ra^-/-^* (lower two rows) larvae at early post-amputation time points (0.5 hpa, 1 hpa, 3 hpa, 6 hpa). Numbers in lower left corners of each image indicates the portion of larvae that showed identical/similar staining to the one shown. scalebar = 100 µm. **B.** Schematic depiction of the mutations in *junba^-/-^* and *junbb^-/-^* fish. **C.** Illustration of the fin fold amputation model in larval regeneration. D. Bright field images of WT, *junba^-/-^, junbb^-/-^ junb^-/-^* DM (double mutant) and *il11ra^-/-^* larvae at different timepoints (24-, 48- & 72 hpa) after fin fold amputations. Red dotted lines indicate the amputation sites while black dotted lines indicate the outline of the regenerated tissue. **E**. Quantification of regenerated area (D, area between red dotted lines and black dotted lines) (24 hpa no significant differences. 48 hpa WT v. *junba^-/-^*: n.s; WT v. *junbb^-/-^*: *** p<0.001; WT v. *junb^-/-^*: *** p<0.001; WT v *il11ra^-/-^*: ***p<0.001. 72 hpa WT v *junba^-/-^*: **p<0.01; WT v *junbb^-/-^*: **p<0.01; WT v *junb^-/-^*: ***p<0.001; WT v *il11ra^-/-^*: ***p<0.001. **F.** Gene set enrichment analysis (GSEA) was performed on the dataset obtained 2 hours after fin-fold amputation in *junb* ^-/-^ vs. WT larvae. The fgsea package from R and REACTOME was used for pathway analysis. The pathways in the representative image were chosen with a p value of <0.02 and an FDR of < 0.1. Transcriptional response to injury 2 hours after fin-fold amputation in *junb* ^-/-^ vs. WT larvae. **G.** Volcano plot of Log2 fold change in expression of *junb ^-/-^*DM vs. WT larvae. A threshold of |log2FC| > 0.585 and p-value < 0.05 was used for the representation, unregulated genes are represented in gray.**H.** Heatmap of genes involved in inflammation, specifically in the *Nfkb* and *Tnf* signalling pathways is represented as a function of z-score, weighted for p-value of each gene.

We next asked which signaling branches downstream of Il-11 contribute to *junb* activation. To this end, we treated wild-type larvae with pharmacological inhibitors targeting canonical and noncanonical pathways downstream of the Il-11 receptor complex (suppl. Fig. 2A). Inhibition of Jak1/Jak2 with Ruxolitinib strongly reduced injury-induced expression of both *junba* and *junbb* (suppl. Fig. 2C). Similarly, pharmacological inhibition of Stat3, a key transcriptional effector downstream of Il-11 receptor signaling, diminished expression of both paralogues (suppl. Fig. 2B). In contrast, inhibition of PI3K signaling with PI-103 did not significantly affect *junba* or *junbb* expression (suppl. Fig. 2C). Notably, inhibition of MEK using PD98059 reduced expression of both *junba* and *junbb* to an extent comparable to that observed upon blockade of Jak/Stat signaling (suppl. Fig. 2C). Together, these data place *junba* and *junbb* downstream of Il-11-dependent Jak/Stat3 and MEK activity during the early injury response.

### Loss of junba and junbb impairs regeneration and shifts the injury response toward inflammatory gene activation

To examine the functional requirement for Junb during regeneration, we generated CRISPR/Cas9-induced mutant alleles for *junba* and *junbb* (Fig. 2B). In both genes, the introduced mutations are predicted to cause frameshifts and premature termination before the bZIP DNA-binding and leucine zipper dimerization domains, consistent with severe loss-of-function alleles (Fig. 2B). We then performed fin fold amputations at 3 days post-fertilization and compared regeneration in wild-type larvae, *junba* and *junbb* single mutants, *junba*; *junbb* double mutants, and *il11ra* mutants (Fig. 2C). Morphological analysis at 24, 48, and 72 hpa revealed impaired regenerative outgrowth in *junb* mutants, with *junbb* mutants displaying a stronger phenotype than *junba* mutants (Fig. 2D, E). Although no significant differences in regenerated area were detected at 24 hpa, defects became evident at 48 and 72 hpa. Quantification of regenerated fin area confirmed a significant reduction in *junbb* mutants and *il11ra* mutants, whereas *junba* mutants showed a milder defect (Fig. 2E). Notably, *junba*;*junbb* double mutants (henceforth referred to as *junb^-/-^* DM) exhibited a severe impairment of regenerative outgrowth, including a pronounced loss of outgrowth in the central fin fold region that resulted in a characteristic V-shaped morphology (Fig. 2D, E). These findings indicate that both *junb* paralogues contribute to fin fold regeneration, consistent with partially redundant functions and a prominent requirement for *junbb*.

The impaired regenerative outgrowth in *junb^-/-^*DM prompted us to investigate whether loss of Junb function alters the early injury-induced transcriptional response. We therefore performed bulk RNA sequencing on tissue collected from wild-type and *junb^-/-^* DM larvae 2 hours after fin fold amputation (Fig. 2F-H). This early time point was chosen to capture transcriptional changes preceding overt morphological differences in regenerate outgrowth. Gene set enrichment analysis of ranked differentially expressed genes revealed positive enrichment of inflammatory pathways in *junb^-/-^*DM, including Toll-like receptor cascades, Interleukin-1 signaling, NF-κB signaling, and TNF-associated pathways (Fig. 2F). This profile was reminiscent of the inflammatory response observed in injured mouse hearts at 24 hours post-injury (Fig. 1B). Consistently, volcano plot analysis showed increased expression of multiple inflammation-associated mediators and signaling components in *junb^-/-^* DM compared with wild-type larvae (Fig. 2G). A weighted z-score heatmap further confirmed coordinated upregulation of Nfκb- and TNF-associated genes in the absence of Junb function (Fig. 2H). Thus, loss of Junb function leads to coordinated early upregulation of immune and inflammatory gene programs after injury. Together with the impaired regeneration phenotype, these data identify Junb as a downstream component of Il-11-dependent injury signaling that constrains early inflammatory gene activation and contributes to maintaining a regeneration-permissive inflammatory state after tissue damage.

### Loss of Junb amplifies inflammatory marker expression and neutrophil recruitment after injury

The early transcriptional activation of inflammatory pathways in *junb^-/-^*DM prompted us to validate whether this dysregulation is reflected at the level of selected inflammatory markers and immune cell recruitment. Because successful regeneration requires a tightly regulated inflammatory response (17,19), we first analyzed the expression of the injury-associated inflammatory markers *il-1β* and *nfκb* after fin fold amputation. In wild-type larvae, both transcripts were induced at 6 hpa, consistent with the transient inflammatory response that accompanies tissue repair and regeneration (Fig. 3A) (18,19). By 24 hpa, *il-1β* expression remained modestly elevated, whereas *nfκb* expression had largely returned toward baseline levels (Fig. 3A) (4).

**Figure 3:**
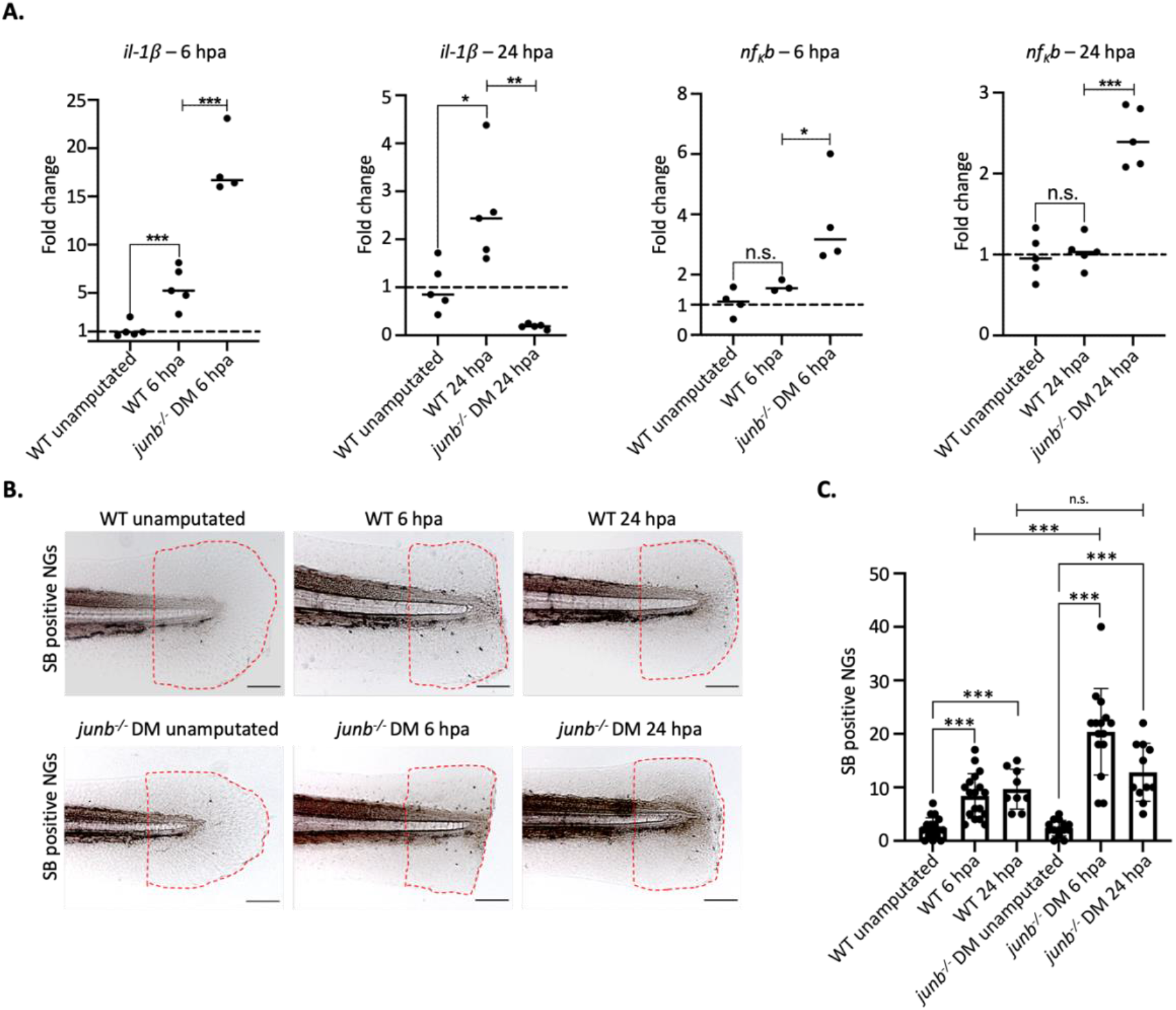
*junb^-/-^*larvae show higher levels of inflammatory markers and higher tissue infiltration of neutrophil granulocytes (=NGs) after fin fold amputation compared to WTs. **A.** RT-qPCR Analysis of the expression of the mRNA for the inflammation associated genes *il1β* and *nfκb* in Wildtype, *il11ra^-/-^*and *junb*^-/-^ DM mutant larvae at 6 and 24 hpa. (n = 5 sample from tissue pooled of 20 larvae, *junb*^-/-^ DM 6 hpa n = 4 samples from tissue pooled of 20 larvae.) Significance is indicated as *p < 0.05; **p < 0.01; ***p < 0.001. **B.** Staining of neutrophil infiltration into the post amputation fin fold tissue of WT and *junb^-/-^* DM larvae at 6 -and 24 hpa via Sudan Black Staining. Red dotted lines mark the area in which neutrophil granulocytes were quantified. Scalebar = 100 µm. **C.** Quantification of counted SB positive neutrophil granulocytes in the post-amputation fin fold area.

In contrast, Junb-deficient larvae displayed markedly enhanced expression of both inflammatory markers at 6 hpa when compared with wild-type larvae (Fig. 3A). At 24 hpa, the response remained dysregulated: *il-1β* expression declined sharply, whereas *nfκb* expression remained elevated (Fig. 3A). Thus, loss of Junb function does not simply prolong a normal inflammatory response but alters the magnitude and timing of inflammatory marker expression after tissue damage.

To determine whether these molecular changes were accompanied by altered immune cell recruitment, we analyzed neutrophil accumulation after fin fold amputation using Sudan Black B staining (Fig. 3B, C). In wild-type larvae, neutrophil numbers increased from approximately two cells in uninjured fins to approximately seven cells at 6 hpa and ten cells at 24 hpa (Fig. 3B, C). Junb-deficient larvae showed substantially increased neutrophil accumulation, with approximately 19 cells detected at 6 hpa and persistently elevated numbers at 24 hpa (Fig. 3B, C). In line with this cellular phenotype, RNA-seq analysis (Fig. 2H) revealed increased expression of *cxcl18b*, a Cxcl-family chemokine previously implicated in neutrophil recruitment (20).

Together, these findings validate the transcriptomic evidence for a hyperinflammatory injury response in Junb-deficient larvae and show that loss of Junb amplifies early inflammatory marker expression and neutrophil recruitment after fin fold amputation.

### Junb deficiency impairs regenerative program activation and induces fibroinflammation

Having established that Junb-deficient larvae mount a hyperinflammatory response after injury, we next asked whether this dysregulation is accompanied by defects in downstream regenerative programs. Inflammation is required for tissue repair, but excessive or mistimed inflammatory activity can interfere with the proliferative and patterning responses needed for successful regeneration. We therefore first assessed cell proliferation after fin fold amputation by EdU incorporation at 24, 48, and 72 hpa (Fig. 4A, B).

**Figure 4:**
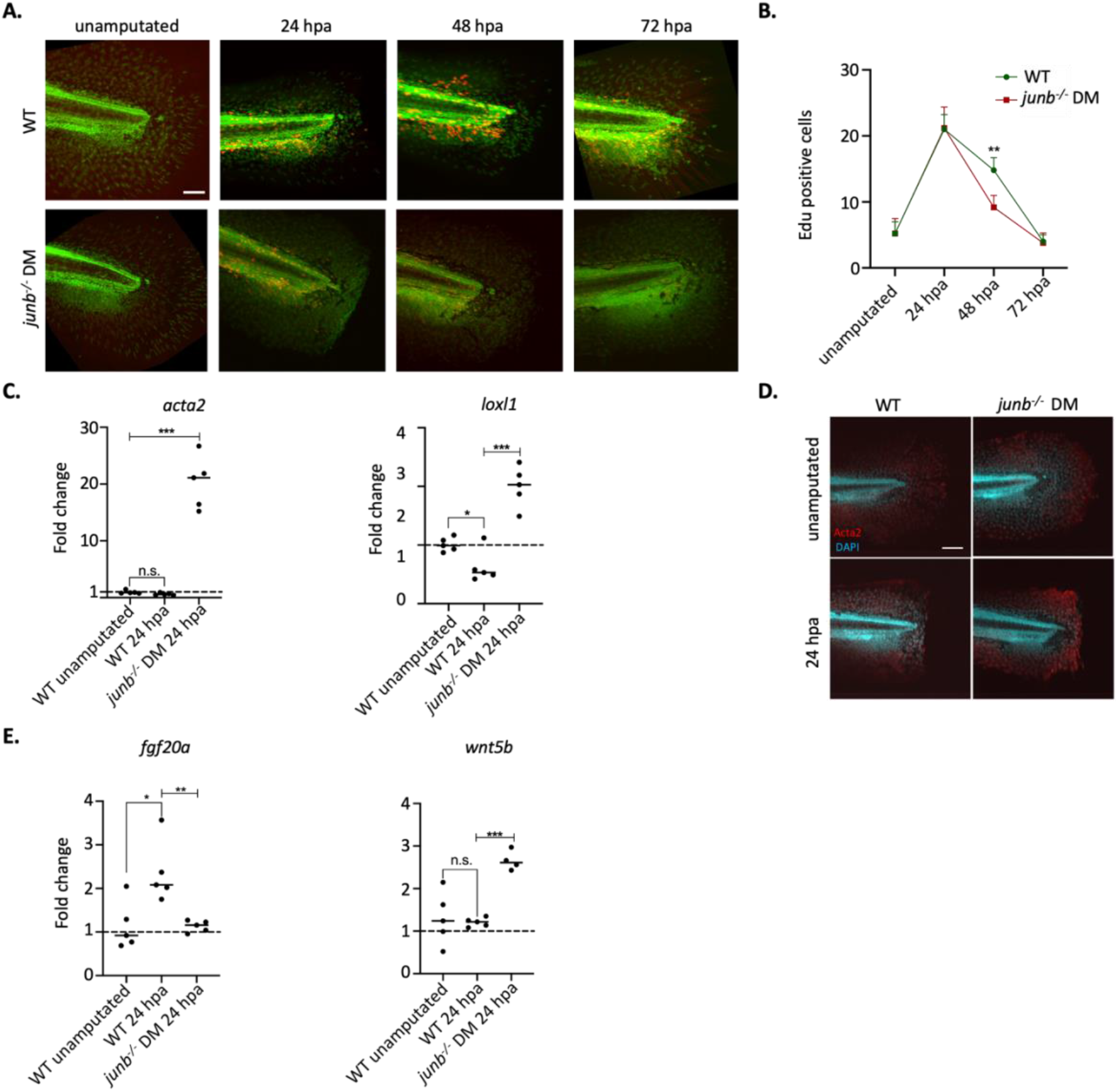
Comparing proliferation, fibrosis and connection to regeneration associated pathways in *junb^-/-^* DM with WT. **A.** Whole mount fluorescence staining of proliferating cells (Edu-Red) and nuclei (green) in WT and *junb^-/-^* larvae unamputated, 24 hpa 48 hpa and 72 hpa. Scalebar = 100 µm. **B.** Quantification of counted Edu positive cells and comparison between WT & *junb^-/-^* DM. Significance indicated by **p < 0.01. **C.** RT-qPCR analysis of the expression of mRNA for the fibrosis associated genes *acta2* and *loxl* in WT, *il11ra^-/-^*and *junb^-/-^* DM larvae at 24 hpa (n = 5 samples containing the pooled tissue of 20 larvae). Significance is indicated as ***p < 0.001. **D.** Whole-mount Immunostaining for Acta2 (red) and DAPI (blue) on WT and *junb^-/-^* at 24 hpa. Scalebar = 100 µm. **E.** RT-qPCR analysis of the expression of mRNA of the genes *fgf20*a and *wnt5b* in WT, *il11ra^-/-^* and *junb^-/-^* DM larvae at 24 hpa (n = 5 samples containing the pooled tissue of 20 larvae). Significance is indicated as *p < 0.05; **p < 0.01; ***p < 0.001.

At 24 hpa, EdU-positive cell numbers were comparable between wild-type and Junb-deficient larvae, consistent with this early phase being dominated by wound closure and inflammatory activation (Fig. 4A, B). By 48 hpa, however, Junb-deficient larvae displayed a substantial reduction in EdU-positive cells in the regenerating fin fold area, particularly near the amputation plane, where proliferative outgrowth normally occurs during larval fin fold regeneration (Fig. 4A, B). By 48 hpa, however, Junb-deficient larvae displayed a substantial reduction in EdU-positive cells in the regenerating fin fold area, particularly near the amputation plane, where proliferative outgrowth normally occurs during larval fin fold regeneration (Fig. 4A, B) (14,19). At 72 hpa, proliferation had declined in both wild-type and mutant larvae, and no significant difference was observed (Fig. 4A, B). These data indicate that loss of Junb impairs proliferation affecting blastema expansion during a defined regenerative window between 24 and 48 hpa resulting in impaired regenerative outgrowth.

We next investigated whether the hyperinflammatory response in Junb-deficient larvae was associated with fibroinflammatory marker expression. RT-qPCR analysis revealed increased expression of *acta2*, a marker associated with myofibroblast activation, in Junb-deficient larvae at 24 hpa (Fig. 4C). Similarly, expression of the extracellular matrix-modifying enzyme *loxl1*, which is associated with matrix crosslinking and fibrotic remodeling, was significantly upregulated (Fig. 4C). In line with these transcriptional changes, whole-mount immunostaining showed enhanced Acta2 presence in mutant fins, most prominently in the mesenchymal compartment (Fig. 4D). Together, these data indicate that loss of Junb promotes early fibroinflammatory marker expression after tissue damage.

Consistent with impaired regenerative outgrowth, Junb-deficient larvae also showed altered expression of regeneration-associated signaling genes. Expression of *fgf20a*, a critical regulator of blastema formation and fin regeneration (21,22), was reduced in mutant larvae compared with wild-type controls at 24 hpa (Fig. 4E). Conversely, *wnt5b* expression was increased in Junb-deficient larvae (Fig. 4E). Since elevated *wnt5b* activity has been linked to impaired blastemal proliferation and reduced fin regeneration, these data suggest that loss of Junb is associated with an imbalance in signaling pathways required for regenerative outgrowth (23).

Altogether, these findings position Junb as an integrator of inflammatory modulation and regenerative activation. Loss of *junb* leads to an impaired proliferative response, disrupted *Fgf* and *Wnt* signaling, and premature expression of fibrotic markers. The convergence of these effects identifies Junb as a molecular switch governing the balance between regeneration and fibrotic repair. These findings extend our understanding of the transcriptional mechanisms underlying regenerative failure and support a broader model in which Junb*/*AP-1 dysregulation disrupts the cellular and molecular framework required for functional tissue restoration.

### AP-1 inhibition supports loss of AP-1 function in Junb-deficient larvae

Junb functions as a component of AP-1 transcription factor complexes, which are rapidly activated downstream of tissue injury and regulate genes involved in stress responses, cytokine signaling, and inflammation (24,25). However, loss of Junb does not necessarily result in a simple loss of AP-1 activity. Instead, altered AP-1 complex composition in the absence of Junb could redirect AP-1-dependent transcription toward an aberrant, neomorphic inflammatory state potentially resulting in excessive or prolonged inflammation, which is often associated with various pathological conditions, including chronic inflammatory diseases and tissue fibrosis (26). To distinguish between these possibilities, we pharmacologically inhibited AP-1-dependent transcription using SR11302, a synthetic retinoid and general AP-1 inhibitor (Fig. 5A).

**Figure 5:**
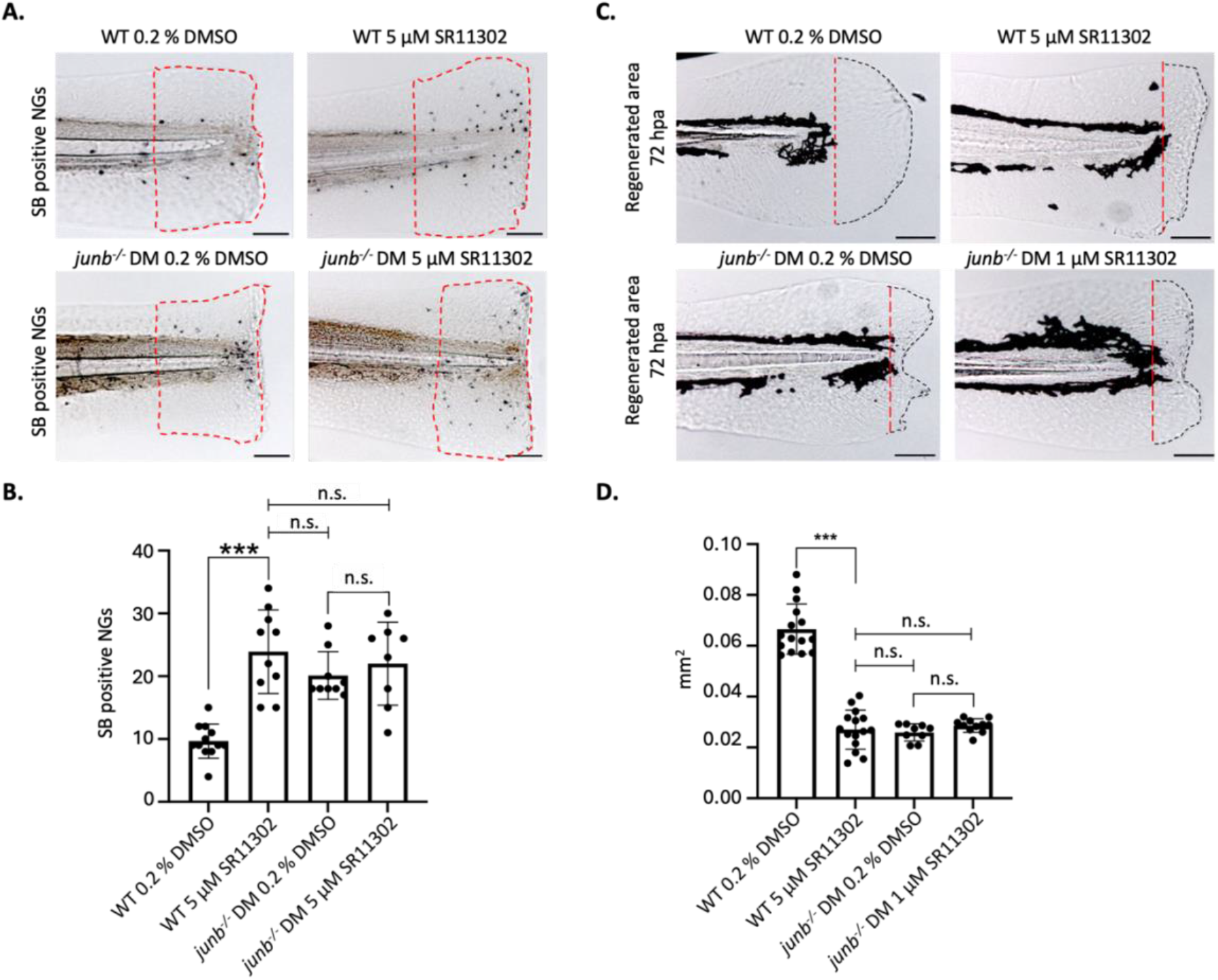
Pharmacological inhibition of the AP-1 complex with the synthetic retinoid SR11302 mimics the *junb^-/-^* associated hyperinflammatory non-regenerative phenotype in WT larvae. **A.** Staining of neutrophil infiltration in the post amputation fin fold area of WT and *junb^-/-^* DM at 6 hpa after treatment with the synthetic retinoid and AP-1 inhibitor SR11302 via Sudan Black B staining. Red dotted lines mark the area in which neutrophil granulocytes were quantified. Scalebar = 100 µm. **B.** Quantification of counted SBB positive neutrophil granulocytes (A) in the post amputation fin fold area. Significance is indicated by ***p < 0.001. **C.** Bright field images of larval fin fold regeneration in WT and junb^-/-^ DM at 72 hpa after treatment with the AP-1 Inhibitor SR11302. Red dotted line indicates the amputation site. Black dotted lines indicate the outline of the regenerated finfold. Area between the red and black dotted lines are measured as regenerated area. Scalebar = 100 µm **D.** Quantification of the regenerated area in WT and *junb^-/-^* DM larvae at 72 hpa after treatment with AP-1 Inhibitor SR11302. Significance is indicated by ***p <0.001.

At 6 hpa, SR11302 treatment increased the number of Sudan Black B-positive neutrophils in wild-type larvae when compared with untreated controls (Fig. 5A, B). Notably, neutrophil accumulation in SR11302-treated wild-type larvae reached levels comparable to those observed in untreated Junb-deficient larvae (Fig. 5A, B). SR11302 treatment did not attenuate the elevated neutrophil accumulation in Junb-deficient larvae; instead, neutrophil numbers remained comparable to those observed in untreated mutants and inhibitor-treated wild-type larvae (Fig. 5A, B). These data indicate that pharmacological AP-1 inhibition is sufficient to increase inflammatory cell recruitment after injury and does not rescue the hyperinflammatory phenotype caused by loss of Junb.

We next assessed whether AP-1 inhibition also affects regenerative outgrowth. Quantification of regenerated fin fold area at 72 hpa showed that SR11302 strongly reduced regenerative outgrowth in wild-type larvae (Fig. 5C, D). Notably, SR11302-treated wild-type larvae also displayed a characteristic V-shaped morphology with reduced central fin fold outgrowth, strongly resembling the phenotype observed in Junb-deficient larvae (Fig. 5C, D). Thus, pharmacological inhibition of AP-1-dependent transcription phenocopies key aspects of Junb loss, including excessive neutrophil recruitment and impaired regenerative outgrowth. Together, these findings support the notion that Junb deficiency primarily compromises an AP-1-dependent function required to restrain inflammatory cell recruitment and support fin fold regeneration. The failure of SR11302 to suppress neutrophil accumulation in Junb-deficient larvae argues against the hyperinflammatory phenotype being driven mainly by neomorphic AP-1 activity from alternative complex composition.

### Anti-inflammatory treatment restores regeneration despite loss of Junb/AP-1 function

In *junb* deficient zebrafish larvae, fin fold amputation triggers hyperinflammation characterized by elevated pro-inflammatory cytokines such as Il-1β, high NF-κB signaling and increased neutrophil recruitment. To test whether hyperinflammation is directly responsible for a reduced regenerative outcome, we investigated the effects of Dexamethasone, a steroidal anti-inflammatory drug (SAID) with potent immunosuppressive properties (27), and ibuprofen, a non-steroidal anti-inflammatory drug (NSAID) that inhibits cyclooxygenase (COX) enzymes to reduce prostaglandin production (28) on *junb* larvae. These two drugs were chosen to target distinct inflammatory pathways, dexamethasone acting upstream of cytokine production and inflammatory gene expression at the transcriptional level, and ibuprofen acting downstream on eicosanoid production. Both drugs were assessed for their ability to modulate the inflammatory response and improve fin regeneration in *junb^-/-^* DM mutants.

Dexamethasone treatment for 6 h strongly reduced the number of Sudan Black B-positive neutrophils at the amputation site in Junb-deficient larvae from 19 +/- 3 to 9 +/- 2 cells, bringing neutrophil numbers close to wild-type levels (7 +/- 2 cells; Fig. 6A, B). By contrast, dexamethasone had little effect on neutrophil numbers in wild-type larvae (7 +/- 2 versus 6 +/-2 cells; Fig. 6A, B). We next determined whether dampening this early inflammatory response affected regenerative outgrowth. At 72 hpa, Dexamethasone did not reduce fin fold regeneration in wild-type larvae (100 +/- 7% versus 98 +/- 8%; Fig. 6 C, D). In Junb-deficient larvae, however, dexamethasone substantially improved regenerative outgrowth in the central blastema region from 55 +/- 10% to 70 +/- 9% (Fig. 6C, D). The rescue was incomplete, as dexamethasone-treated mutants still regenerated less efficiently than wild-type controls (Fig. 6C, D). Thus, reducing inflammatory activation by dexamethasone improves, but does not fully normalize, the regenerative response in Junb-deficient larvae.

**Figure 6:**
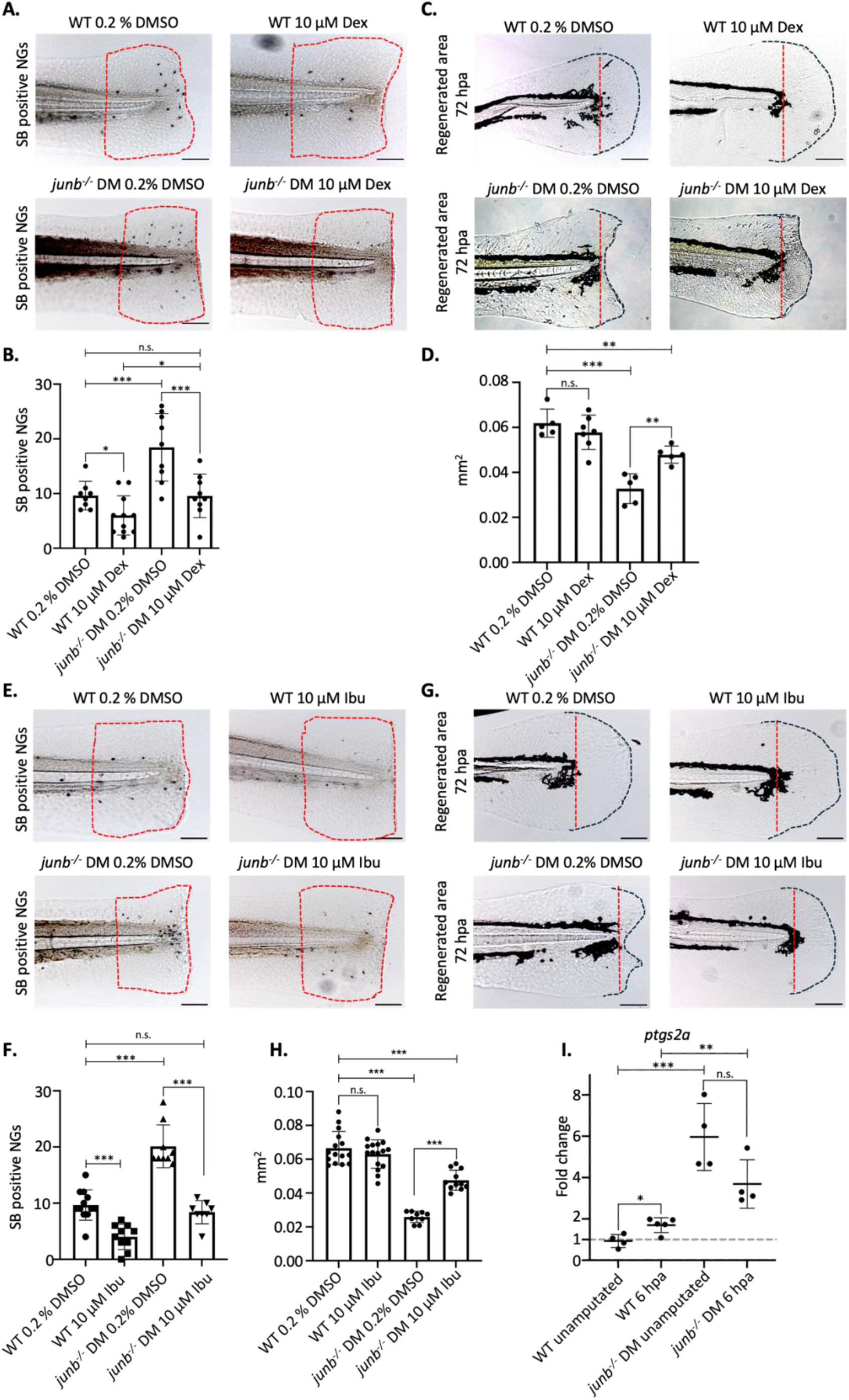
Steroidal and non-steroidal modulation of hyperinflammation partially restores regeneration in the *junb^-/-^* DM. **A.** Staining of neutrophil infiltration in WT and *junb^-/-^*DM larvae at 6 hpa after treatment with the steroidal anti-inflammatory drug (SAID) Dexamethasone (Dex) via Sudan Black B Staining. Red dotted outlines indicate the area in which neutrophil granulocytes were counted. Scalebar = 100 µm **B.** Quantification of counted SB positive neutrophil granulocytes (A) in the post amputation fin fold area. Significance is indicated by *p < 0.05, *** p < 0.001**. C.** Bright field images of larval fin fold regeneration in WT and *junb^-/-^* DM at 72 hpa after treatment with the the SAID Dexamethasone. Red dotted line indicates the amputation site. Black dotted lines indicate the outline of the regenerated fin fold. Area between the red and black dotted lines are measured as regenerated area. Scalebar = 100 µm **D.** Quantification of the regenerated area in WT and *junb^-/-^*DM larvae at 72 hpa after treatment with Dexamethasone. Significance is indicated by ***p < 0.001. **E.** Staining of neutrophil infiltration in WT and *junb^-/-^*DM larvae at 6 hpa after treatment with the non-steroidal anti-inflammatory-drug (NSAID) Ibuprofen (Ibu) via Sudan Black B Staining. Red dotted outlines indicate the area in which neutrophil granulocytes were counted. Scalebar = 100 µm. **F.** Quantification of counted SBB positive neutrophil granulocytes (E) in the post amputation fin fold area. Significance is indicated by *** p < 0.001**. G.** Bright field images of larval fin fold regeneration in WT and *junb^-/-^* DM at 72 hpa after treatment with Ibuprofen. Red dotted line indicates the amputation site. Black dotted lines indicate the outline of the regenerated fin fold. Area between the red and black dotted lines are measured as regenerated area. Scalebar = 100 µm. **H.** Quantification of the regenerated area in WT and *junb^-/-^*DM larvae at 72 hpa after treatment with Ibuprofen. Significance is indicated by ***p < 0.001. **I.** RT-qPCR analysis of the expression of mRNA for the prostaglandin synthesis *ptgs2a* in WT, and *junb^-/-^* DM larvae at 6 hpa (WT unamputated and WT 6 hpa n = 5 samples containing the pooled tissue of 20 larvae. *junb^-/-^*DM unamputated and *junb^-/-^*DM 6 hpa n = 4 samples containing the pooled tissue of 20 larvae). Significance is indicated as *p < 0.05, **p < 0.01, ***p < 0.001. **J.** RT-qPCR analysis of the expression of mRNA for the prostaglandin degrading enzyme *ptgr1* in WT, and *junb^-/-^* DM larvae at 6 hpa (WT unamputated and WT 6 hpa n = 5 samples containing the pooled tissue of 20 larvae. *junb^-/-^*DM unamputated and *junb^-/-^* DM 6 hpa n = 4 samples containing the pooled tissue of 20 larvae). Significance is indicated as **p < 0.001.

Since Dexamethasone can also influence blastema marker expression and regenerative signaling pathways (29), we next asked whether a more downstream anti-inflammatory intervention would produce a similar effect. Ibuprofen treatment reduced neutrophil accumulation in Junb-deficient larvae from 19 +/- 3 to 10 +/- 3 cells, while neutrophil numbers in wild-type larvae remained unchanged (7 +/- 2 versus 7 +/- 2 cells; Fig. 6E, F). Importantly, ibuprofen did not impair normal fin fold regeneration in wild-type larvae (100 +/- 7% versus 101 +/- 6%; Fig. 6G, H). In Junb-deficient larvae, however, ibuprofen substantially restored regenerative outgrowth from 55 +/- 10% to 68 +/- 11%, again without fully reaching wild-type levels (Fig. 6G, H). The concordant effects of dexamethasone and ibuprofen indicate that the impaired regeneration of Junb-deficient larvae is functionally linked to excessive inflammatory signaling rather than to a drug-specific effect.

As Ibuprofen acts through cyclooxygenase inhibition, we next examined expression of *ptgs2a*, a key enzyme involved in prostaglandin synthesis. RT-qPCR analysis showed that *ptgs2a* expression was elevated in unamputated Junb-deficient larvae compared with wild-type controls and remained higher in mutants after amputation (Fig. 6I). These data are consistent with increased cyclooxygenase-associated inflammatory signaling in the absence of Junb. Together, these findings identify hyperinflammation as a major functional component of the regeneration defect caused by loss of Junb. Notably, regenerative outgrowth can be substantially restored by steroidal or non-steroidal anti-inflammatory treatment despite the underlying genetic loss of Junb/AP-1 function. Thus, Junb-dependent AP-1 activity does not simply promote regeneration through a linear pro-regenerative program, but is required to maintain an inflammatory state that remains compatible with tissue restoration (30).

## DISCUSSION

Tissue regeneration requires activation of repair programs while avoiding a persistent fibroinflammatory state. This distinction is particularly evident after cardiac injury, where zebrafish regenerate damaged myocardium, whereas adult mammals typically respond with prolonged inflammation, fibrotic remodeling, and scar formation (8). Our comparative analysis indicates that regenerative and non-regenerative injury responses diverge early after damage. Although both zebrafish and mouse hearts activate inflammatory and hemostatic programs, mouse hearts display broad NF-κB/TNF signaling and induction of multiple IL-6 family cytokines, whereas zebrafish hearts show a more restricted response dominated by Il-11 and accompanied by proliferative pathway activation. Importantly, this divergence was not limited to the heart. Across the tested injury contexts, non-regenerating mammalian tissues shared a related inflammatory signature that separated them from regenerative species, independent of tissue identity. Thus, regenerative outcome appears to depend not on the absence of inflammation, but on whether inflammatory signaling is constrained into a regeneration-permissive state.

Our data identify Junb as an Il-11-responsive AP-1 component that contributes to this inflammatory restraint. Previous work established Il-11 signaling as a pro-regenerative pathway in zebrafish, required for tissue restoration and suppression of fibrotic remodeling in multiple organs (7,33). Here, we extend this model by showing that Il-11-associated Jak/Stat3 and MAPK signaling promotes rapid induction of *junba* and *junbb* after injury. This places *junb* within the immediate early injury response and links Il-11 signaling to transcriptional control of inflammation. Thus, Il-11-dependent regeneration is not only associated with later tissue remodeling and scar suppression, but also with early regulation of the inflammatory state that follows damage.

Loss of Junb shifts this response toward a hyperinflammatory and fibroinflammatory phenotype. Junb-deficient larvae show impaired regenerative outgrowth, increased inflammatory pathway activation, enhanced neutrophil recruitment, reduced proliferation during the outgrowth phase, and induction of fibroinflammatory markers. Notably, this phenotype resembles key aspects of the non-regenerative mammalian injury signature, including increased NF-κB/TNF and interleukin signaling. These findings support a model in which Junb prevents a normally regenerative zebrafish injury response from entering a mammalian-like inflammatory state. In this view, regenerative and non-regenerative outcomes are not defined solely by the presence or absence of regenerative programs, but by whether early injury-induced inflammation remains within a permissive range (34). The AP-1 inhibitor experiments support this interpretation. Because Junb functions within AP-1 complexes, loss of Junb could either reduce AP-1-dependent activity or redirect AP-1 signaling through altered complex composition. Pharmacological AP-1 inhibition phenocopied central aspects of Junb deficiency in wild-type larvae, including increased neutrophil accumulation and impaired regenerative outgrowth, and did not suppress the hyperinflammatory phenotype in larvae mutant for *junba* and *junbb*. These data are consistent with a model in which Junb deficiency primarily reflects loss of an AP-1-dependent function required to restrain inflammation and support regeneration.

The anti-inflammatory rescue experiments provide the strongest functional support for this model. Dexamethasone and Ibuprofen reduced excessive neutrophil recruitment and substantially restored regenerative outgrowth in Junb-deficient larvae, without impairing regeneration in wild-type larvae. This is striking because the underlying genetic loss of Junb/AP-1 function remains present. Thus, the regeneration defect is not explained solely by the absence of a pro-regenerative transcription factor. Rather, a major component of the phenotype arises from the inflammatory state generated in the absence of Junb/AP-1 function. The incomplete rescue further suggests that Junb/AP-1 also contributes to additional injury-induced transcriptional programs, but the ability of two mechanistically distinct anti-inflammatory treatments to improve regeneration demonstrates that inflammatory balance is a functional determinant of regenerative outcome.

These findings have broader implications for regenerative biology. AP-1 motifs are enriched in regeneration-responsive enhancers, and evolutionary changes in these elements have been proposed to separate regenerative from non-regenerative injury responses (35). Our data suggest that AP-1 activity should not be viewed as a generic injury signal. Instead, specific AP-1 components such as Junb may determine whether injury-induced transcription is resolved into a regenerative or fibroinflammatory state. It will therefore be important to define the direct targets and cellular sites of Junb function, and to test whether Junb similarly constrains inflammation during adult zebrafish heart regeneration.

Together, our study identifies an Il-11-Junb/AP-1 axis that controls the inflammatory quality of the early injury response. Junb limits hyperinflammatory gene activation and neutrophil recruitment, thereby maintaining a tissue environment compatible with proliferative outgrowth and regeneration. Loss of Junb shifts a regenerative zebrafish injury model toward a mammalian-like fibroinflammatory state, whereas anti-inflammatory treatment substantially restores regeneration despite the underlying AP-1 defect. By linking Il-11 signaling to AP-1-dependent inflammatory restraint, these findings provide a mechanistic explanation for how regenerative species may avoid the conserved inflammatory state associated with mammalian repair failure.

## MATERIALS AND METHODS

### Zebrafish husbandry

Adult zebrafish were maintained on a 12h day/12h night cycle. The water temperature was constantly 28.5 °C. Up to 15 fish were kept in a tank with a volume of eight liters. The animals were fed artemia (Sanders Great Salt Lake artemia cysts) between 7:00 a.m. and 12:00 p.m. and between 4:00 p.m. and 7:00 p.m. with dry food (Sera Micron Nature Anzuchtfutter; Sera ImmunPro Mini Nature Growth Food; Dr. Bassleer Biofish food). Different fish lines were used for the experiments. Fish of the AB wild-type line served as a control group. Furthermore, zebrafish with mutated *il11ra*, *junba* and *junbb* were used. Zebrafish husbandry was performed under standard conditions in alignment with ethical and animal welfare guidelines approved by the ethics committee for animal experiments.

### Generation of zebrafish mutant lines

Mutant alleles were generated by injecting Cas9 mRNA (300 pg, from pT3TS-nCas9n, Addgene plasmid 46757) and 50 pg of gRNA for *junba* (5’-GGACGGATTCGTCAAAGCGC-3’) and *junbb* (5’-GGGTTACGGTCACAACGACG-3’) in one-cell stage embryos from the wild-type AB background (gRNA sequences from Kiesow et al, 2015, *Scientific Reports*). The *junba* mutant allele used here is an 8-nucleotide deletion allele leading to a frameshift and premature stop codon (562_569del_fsX51, ENSDARG00000074378), and *junbb* is an insertion/deletion mutant leading to a frameshift and premature stop codon (354_355del353_354insT_fsX12, ENSDARG00000104773).

### Sample collection

At 72 hours post fertilization, larval zebrafish were gently anesthetized using 0.016% tricaine (Cat. 442848; Pharmaq) solution in egg water. Under precise observation using a stereomicroscope, the fin folds situated posterior to the notochord were carefully amputated using a scalpel. Subsequently, the zebrafish were allowed to undergo fin fold regeneration at a controlled temperature of 28°C for the specified durations. For the purpose of gene expression analysis, tissue samples located posterior to the yolk extension were meticulously collected (7). For the homozygous samples (wild type, *junba^-^*^/-^ x *junbb ^-^*^/-^and *il11ra^-/-^*) a total of 15 fins per genotype were placed in a 1.5mL tube on ice. For every experiment, 5 replicates (each containing 15 fins) were used. For sample collection that did not need further genotyping, 20-25 fins were collected for each replicate. The samples were then correctly labeled, excess water was removed and the tubes were stored at -80 °C until the experiment was continued. For analyzing the single mutants (either *junba^-^*^/-^ or *junbb^-^*^/-^) genotyping needed to be performed. Each larva was placed separately under the microscope; fin amputation was performed. The isolated fin was collected and placed in a 0.2 mL tube of an 8-stripe. The rest of the larva was collected and placed in a second 8-stripe 0.2mL tube. The respective tube of fin and corresponding head were labeled with the same number. The larger head part of the larva was further used for genotyping while the fins were stored at -80 °C. After successful genotyping, about 15 larval fins with the same desired genotype were transferred to a 1.5 mL Eppendorf tube to continue further procedure.

### Reverse transcription quantitative polymerase chain reaction

Analysing geneexpression involved isolating total RNA from a pool of 15 dissected larvae using the RNA Clean and Concentrator kit (Zymo Research). Following this, at least 200ng of total RNA underwent reverse transcription using the Maxima First Strand cDNA Synthesis Kit (Thermo Fisher Scientific) as per the manufacturer’s instructions. mRNA levels were normalized against *rpl13a* mRNA levels. All reactions were conducted utilizing the SYBR Green PCR Master Mix (Thermo Fisher Scientific) on the CFX Connect Real-Time System (Bio-Rad) or QuantStudio 3 (applied biosystems, Thermo Fisher Scientific). Primer sequences can be found in Table 1.

**Table 1:**
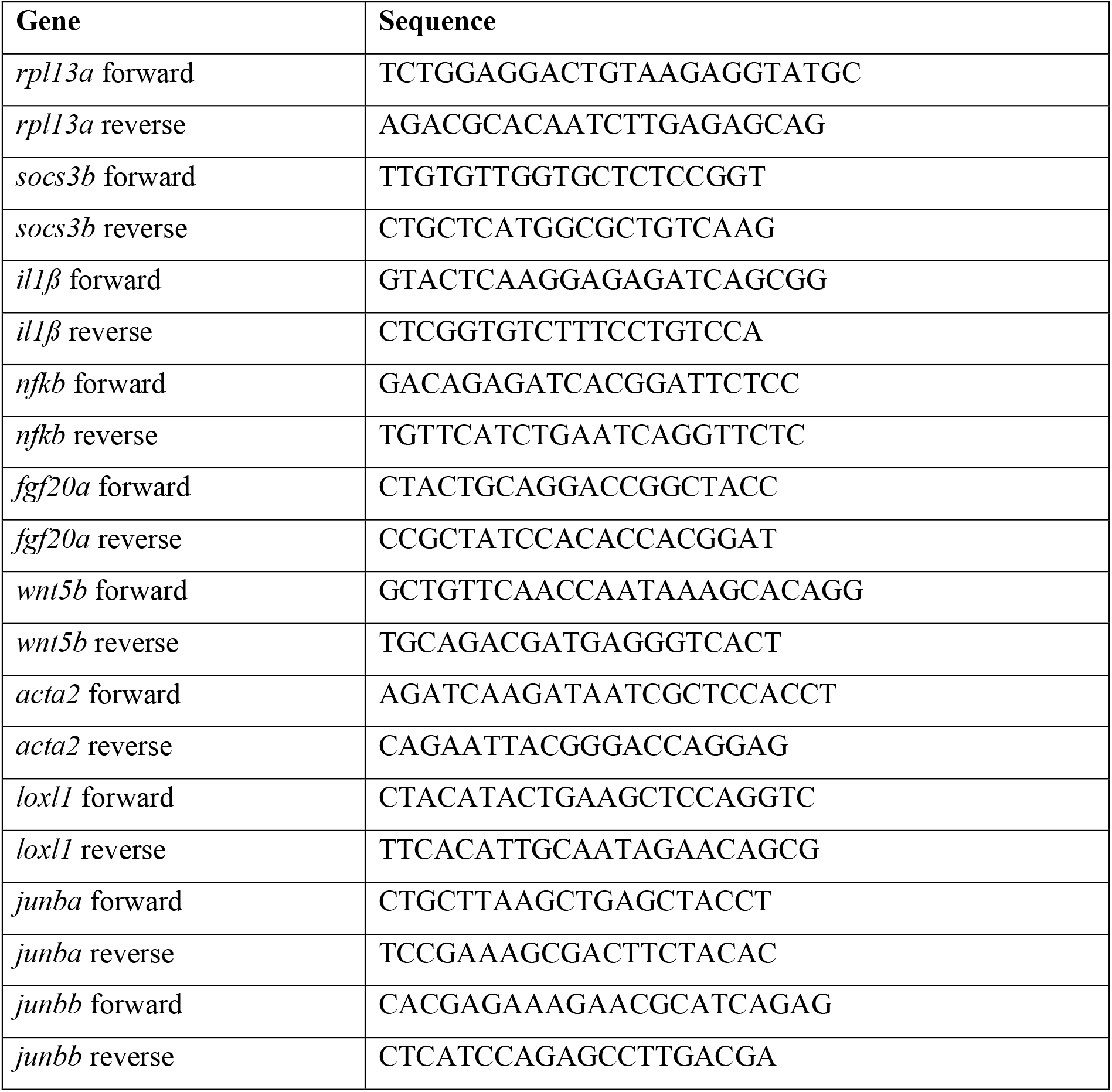
Primer sequences for gene expression analysis.

### Genotyping

#### High Resolution Melt analysis

For genotyping, genomic DNA was isolated by adding 30 μL of a 50 mM NaOH solution to each larva and incubated for 15 minutes at 95 °C in the thermocycler. Then 3 μL of a 1 M Tris solution with a pH value of 8 were added to the mixture, mixed and placed in the centrifuge at 4000 rpm for 1 minute. In a 96-well plate, 5 μL of a 2X master mix SYBR Green, 3.5 μL autoclaved water, 1 μL gDNA solution of the larvae and 0.5 μL of a 10 μM primer mix of the mutation (*junba* or *junbb*) was added. The primer sequences are given in Table 2 in the appendix. After all components had been pipetted, the well-plate was sealed using self-adhesive foil and mixed vigorously using a vortex machine. Melt curve peaks were then analyzed to identify the respective genotype of each sample.

**Table 2:**
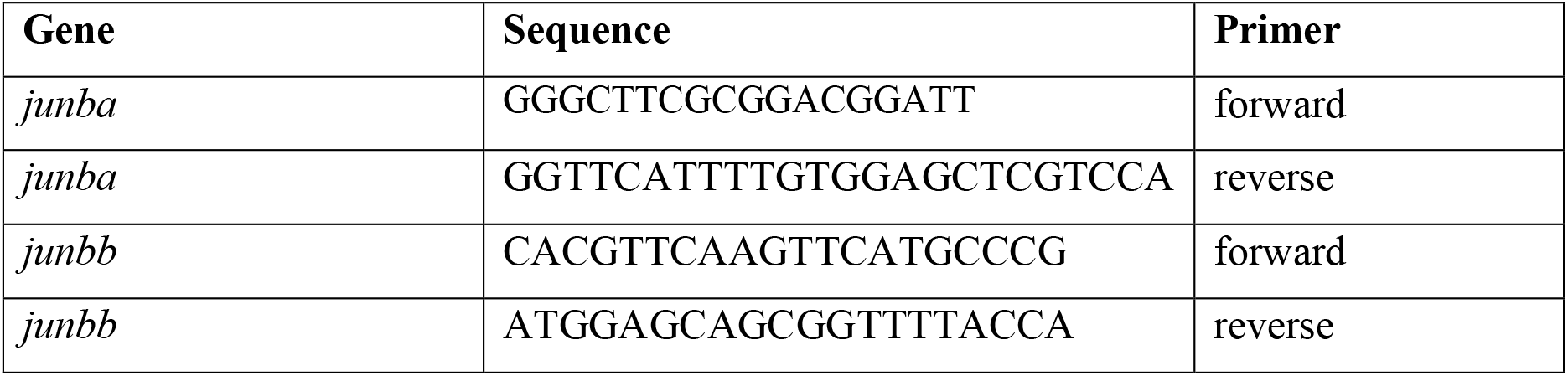
Primer sequence junba and junbb for melt-curve analysis.

### Histological analysis and imaging

#### Lightmicroscopy

After amputation, larvae were fixed in 4% Paraformaldehyde (PFA) at their respective timepoints. After PBTw washes larvae, were transferred through a glycerol series (25%, 50%, 75%) and mounted on glass slides for imaging. Microscopy was carried out with the Leica DM 4000 B Microscope and the attached LEICA DFC 420C and images were documented with the LAS V3.8 Software. Regrowth was measured and Sudan Black positive Neutrophils were counted using FIJI (36) and Quantified using GraphPad Prism V6 /V10.

### EdU staining

Cell proliferation was assessed using the Click-iT™ EdU Imaging Kit (Invitrogen). Zebrafish larvae were incubated in 0.5 mM EdU and 0.5% DMSO in egg water for 6 h at the desired time points following fin fold amputation or in uncut controls (12 larvae/well in 6-well plates). After incubation, larvae were fixed in 4% PFA overnight at 4 °C, washed with PBS/0.1% Tween-20, and permeabilized with Proteinase K (10 μg/mL, 20 min). Following additional washes in PBDT and blocking with 3% BSA, the EdU detection reaction was performed according to the manufacturer’s instructions using Azide 647. Nuclear Green DCS1 (ab138905, Abcam) was used as a counterstain at a dilution of 1:1000. Samples were washed and transferred through a glycerol series (25%, 50%, 75%) for storage at 4 °C in the dark. Larvae were mounted in glycerol and imaged using the Zeiss Axioscop 2 - LSM5 Live or the Zeiss AXIO-LSM5.

### Antibody staining

Fin amputation was performed and larvae were kept under standard conditions for a period of time in accordance with the respective experiment. Embryos were then fixed with 4% PFA overnight. The following day, larvae were washed 5 times in PBS for 10 minutes. Proteinase K (10μg/mL) in 1 mL PBS/0.1% Tween20 was used to permeabilize the embryos. The embryos were kept in Proteinase K for 20 minutes and this process was stopped using 1 mL PBDT to rinse out and further washed 2-3x with PBDT. Blocking was performed by adding 400μL PBDT and 8μL of 4% Goat Serum (Sigma Aldrich) and placing the tube on a shaker for 60 minutes at room temperature. Blocking solution was removed and 200μL PBDT were added. The primary antibody ACTA2 (Sigma F3777) 1:100 in 400μL PBDT were added and the tubes were placed at 4°C overnight on a shaker in the dark. The next day, primary antibody was removed and a quick wash with 1mL PBDT was performed and consecutively 4-5 times washing with PBDT with 30 minutes in between. Secondary antibody 1:500 in 0.5mL PBDT was added, and the tubes were placed on a shaker at 4°C overnight. The following day, procedure continued with 2x quick wash with PBDT and subsequent washing 4x with PBDT every 30 minutes. Embryos were fixed by adding 4% PFA at room temperature for 30 minutes. Fixative was washed out with PBS and a glycerol series with 25%, 50%, 70%,80% was performed and samples were stored at 4°C in the dark.

### Sudan black staining

Larvae were fixed with 4% Methanol-free Paraformaldehyde (Cat.; 28908, Thermo Fisher scientific) in Phosphate buffered saline (PBS) overnight. On the following day Larvae were rinsed and washed with PBS and stained with 0,036% (w/v) Sudan Black B (Cat.: 199664 Sigma Aldrich) in 70 % Ethanol for 1:30 h at room temperature. Followed by the staining larvae are rinsed and extensively washed with 70% Ethanol and progressively rehydrated by PBS + 0,1% Tween20 (PBT). To prevent the larval pigmentation from interfering with the evaluation of stained neutrophil granulocytes, Larvae are depigmented with a depigmentation solution containing 10% KOH and 30% H_2_O_2_ for 10 minutes followed by rinsing with PBT and preparation for Imaging by progressive transfer into different Glycerol concentrations.

### HCR FISH staining

The staining was performed according to the protocol provided by the Manufacturer (Molecular Instruments (37). Furthermore, all the used reagents necessary for the protocol (Probes, Amplifiers, Buffers) were ordered from Molecular Instruments.

### Drug treatment

To monitor the influence of the used drugs on the inflammation response by observing neutrophil granulocyte recruitment, 3 dpf old larvae underwent fin fold amputation and were treated for 6 / 24 hours post amputation (hpa) by adding the drugs in their appropriate concentration to the Egg medium and leaving the larvae for the mentioned time at 28 °C. For the larvae that were used for Sudan Black staining the following concentrations were used: 10 µM dexamethasone (D2915, Sigma Aldrich), 10 µM Ibuprofen (I4883, Sigma Aldrich) 5 µM SR11302 (HY-15870, MedChemExpress). Since the drugs were dissolved and diluted in Dimethylsulfoxid (DMSO) (34869, Sigma Aldrich), 0.2 % DMSO was used on control larvae. After the 6 h treatment larvae were killed with 0.4 % Tricaine (E10521, Sigma-Aldrich) fixed with 4% Paraformaldehyde (043368.9M, ThermoScientific) overnight and used for Sudan black staining subsequently as described above. Regeneration Experiments including with the above mentioned Inhibitors that went over the larvae age threshold of 5 dpf were approved by the comittee of ethics in animal experiments. (Proposal G20/2025).

To observe the regenerative behavior of treated larvae, 3 dpf old larvae underwent fin fold amputation and were treated for 72 hours by adding the drugs in their appropriate concentrations to the egg medium. For Dexamethasone as well as Ibuprofen 10 µM was used for the treatments. SR11302 showed a toxic effect at a concentration 10 µM for both the wild-type as well as the *junb^-/-^*larvae during the prolonged exposition in the regeneration experiment. Subsequent testing revealed that for WT 5 µM and for *junb^-/-^* 1 µM were suitable and therefore the concentrations used in the experiments. Similar to the treatments to observe inflammation response, 0.2 % DMSO was used as a control. After the treatment the larvae were euthanized with 0.4 % Tricaine, fixed with 4% Paraformaldehyde overnight and used for evaluation as described above.

### Mouse heart injury and sample preparation

Mouse heart injury was performed via LAD ligation using adult male mice (10-12 weeks old) following safety and animal experimentation rules as per proposal number B2/2062. Samples were obtained from C57BL/6J mouse mice subjected to myocardial infarction by ligation of the left anterior descending (LAD) coronary artery at a non-proximal position to avoid excessive infarct size and cardiac rupture, resulting in medium-sized infarcts. At the time of sacrifice, the infarct zone was defined as the visibly pale myocardial tissue distal (below) to the ligation site, while the remote zone corresponded to tissue proximal (above) to the ligation; in both cases, only ventricular tissue was collected at 24 hours post-injury, excluding the right ventricle and atria. Control samples consisted exclusively of left ventricular tissue. Total RNA was isolated from individual hearts (one heart per sample, without pooling) using TriPure reagent (11667165001, Roche) followed by DNAseI treatment according to the manufacturer’s protocol.

### Zebrafish cryoinjury and sample preparation

Zebrafish heart injury was performed via cryoinjury in accordance with institutional (Max-Planck-Gesellschaft) and national ethical and animal welfare guidelines approved by the ethics committee for animal experiments at the Regierungspräsidium Darmstadt, Germany. Adult zebrafish (4–8 months post-fertilization) were anesthetized using 0.016% tricaine in system water and positioned ventral side up on a moist sponge. A thoracic incision was performed to expose the heart, after which a liquid nitrogen–precooled cryoprobe was applied to the ventricular apex and held in place until thawing occurred. Following the procedure, fish were transferred to fresh system water to recover. Samples were collected at 24 hours post injury and snap-frozen in Trizol. RNA isolation was performed as per manufacturer’s instructions.

### RNA sequencing data processing and analysis

RNA integrity and library quality were assessed using the LabChip Gx Touch (Perkin Elmer). Input RNA (99–249 ng) was normalized prior to library preparation, and sequencing was performed on an Illumina NextSeq500 platform. Reads were quality-trimmed with Trimmomatic (v0.39) using a sliding window (5 nt, Q15) and minimum length of 15 nt, then aligned to the appropriate Ensembl genome (mm39, release 109) with STAR (v2.7.11b). Alignments were filtered to remove duplicates, multimapping, ribosomal, and mitochondrial reads, and gene-level counts were generated with featureCounts (v2.0.4). Count data were normalized and analyzed for differential expression using DESeq2 (v1.36.0), with significance defined as mean count >5, adjusted p-value <0.05, and |log2FC| >0.585. Annotation was supplemented with UniProt data.

### Analysis and Data representation

#### Gene-set enrichment analysis

Gene set enrichment analysis (GSEA) was performed using the fgsea package in R. Genes were ranked by log2 fold change, and duplicate entries were removed by retaining the value with the highest absolute fold change. Reactome pathway gene sets were obtained from MSigDB via the msigdbr package and used to construct pathway collections. Enrichment analysis was conducted with 50,000 permutations (200,000 for zebrafish larvae), and p-values were adjusted using the Benjamini–Hochberg method. Data was plotted using matplotlib, the cut-off threshold was p-value< 0.05 for all pathway analyses, with details mentioned in the figure captions.

### Heat-map

Heatmaps were generated in Python using pandas, NumPy, matplotlib, and seaborn for mouse and zebrafish heart injury datasets

For visualization, gene expression values were standardized by calculating row-wise Z-scores across samples 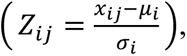, where *x_ij_* is the expression of gene *i*in sample *j*, and *μ*_i_, *σ_i_* are the mean and standard deviation of gene *i*. To incorporate statistical significance, Z-scores were weighted by differential expression p-values using a normalized negative log transformation:

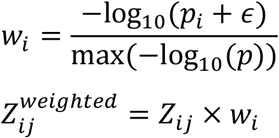

where *p_i_*is the gene-level p-value, *ε* = 10^−10^prevents undefined values, and weights are scaled between 0 and 1. This approach emphasizes genes with stronger statistical support while preserving relative expression patterns. Heatmaps were visualized using a centered blue–white–red colour scale (range −1.5 to 1.5). The code for figure visualization was generated with the help of ChatGPT 5.2.

### Volcano plot

Volcano plots were generated in Python using pandas, NumPy, and matplotlib from datasets for mouse and zebrafish. Gene expression values were filtered for valid numeric entries, and p-values were transformed as −log_10_(*p*). Genes were classified as upregulated or downregulated using thresholds of ∣ log_2_ FC ∣> 0.585and p-value < 0.05. Data from both species were combined into a single plot with species-specific color coding and fold-change–dependent intensity gradients, while non-significant genes were shown in gray. The code for figure visualization was generated with the help of ChatGPT 5.2.

### Dataset analysis

Datasets representing regenerative ((GSE126701, GSE145497, GSE106269) and non-regenerative injury contexts (GSE130438, GSE305817) were analysed for regenerative scores (GO:0042246) and inflammatory signalling as indicated. Differential injury responses were analyzed within each dataset separately. For bulk RNA-seq and processed count tables, published expression matrices were imported and gene-level injury-associated log2 fold changes were calculated using the selected injury and control groups. For the larval zebrafish tail and axolotl limb single-cell datasets, descriptive pseudo-bulk profiles were generated by aggregating control and injury-state cells and calculating mean injury-versus-control log2 expression differences. Cross-species comparison was performed on within-dataset effect sizes rather than pooled expression values. The code was generated with the help of GPT-5.3-Codex.

### Statistics

At least three independent tests were carried out. From three independent tests, the statistical evaluation was carried out using the *GraphPad Prism 6 Software.* The *students’ t test* was used. Here, p < 0.05 was defined as statistically relevant. Data is presented as single points with mean bars or as mean bars plus standard deviation. One star “*” refers to a statistical significance of p < 0.05, two stars “**” refer to a statistical significancy of p < 0.01 and three stars “***” refer to a statistical significance of p < 0.001.

## Acknowledgments

We thank M. Rana, N. Ahmed, L. M. Rafi, C. Troidl, and K. Preissner for comments on the manuscript. We also thank our animal staff for excellent support.

## Funding

This work was supported by the DFG CRC 1213, project B01 and project 496770602 (both SR), as well as the Excellence Cluster Cardio-Pulmonary Institute (SR).

## Author contributions

Conceptualization: JI, EB, SA and SR

Methodology: JI, EB and SR

Investigation: JI, EB, AM

Bioinformatics Data Analysis: AM, SG

Provided Resources: LC-E, JP, SG, DYRS, STS, AB, SR

Writing: JI, EB, AM, SR

With comments and suggestions from all the authors.

Review and editing: SA, JP, DYRS, AB, SR

Supervision: SR

Project administration and funding acquisition: SR

## Competing interests

The authors declare that they have no competing interests.

## Data and materials availability

All data needed to evaluate the conclusions in the paper are present in the paper and/or the Supplementary Materials.

Newly compiled RNA-seq data are deposited to Gene Expression Omnibus (GEO) (https://www.ncbi.nlm.nih.gov/geo/query/acc.cgi?acc=GSE339391) currently under Embargo for reviewers only (reviewer token: qvqtkwemptmrxoj) but will be made publicly available after acceptance.

---

**Supplementary Figure 1:**
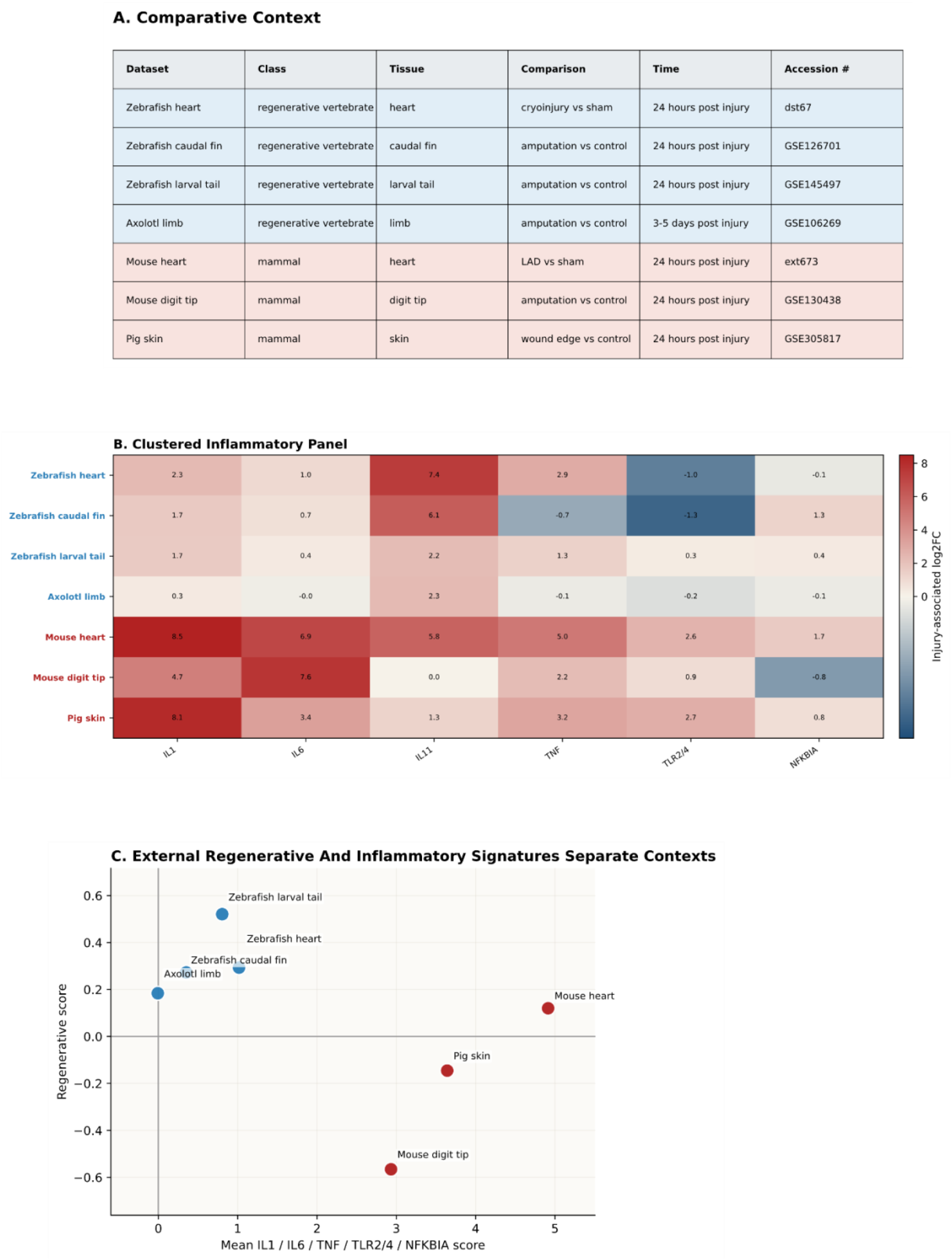
Comparison between regenerative vs. non-regenerative vertebrates after injury. **A.** List of datasets used for a curated cross-context comparison. Datasets include regenerative vertebrate injuries in zebrafish heart, zebrafish caudal fin, larval zebrafish, axolotl, together with non-regenerative mammalian injuries in mouse heart, mouse digit tip, and pig skin **B.** Graphical visualization of inflammatory panel across datasets after hierarchical clustering based on log2 fold change values. **C.** Scatter plot of canonical inflammatory-node score (‘IL1’, ‘IL6’, ‘TNF’, ‘TLR2/4’, and ‘NFKBIÀ) against a curated regenerative score. The regenerative axis is derived from ‘GOBP_TISSUE_REGENERATION’ (‘GO:0042246’), retrieved from MSigDB mouse collections.

**Supplementary Figure 2:**
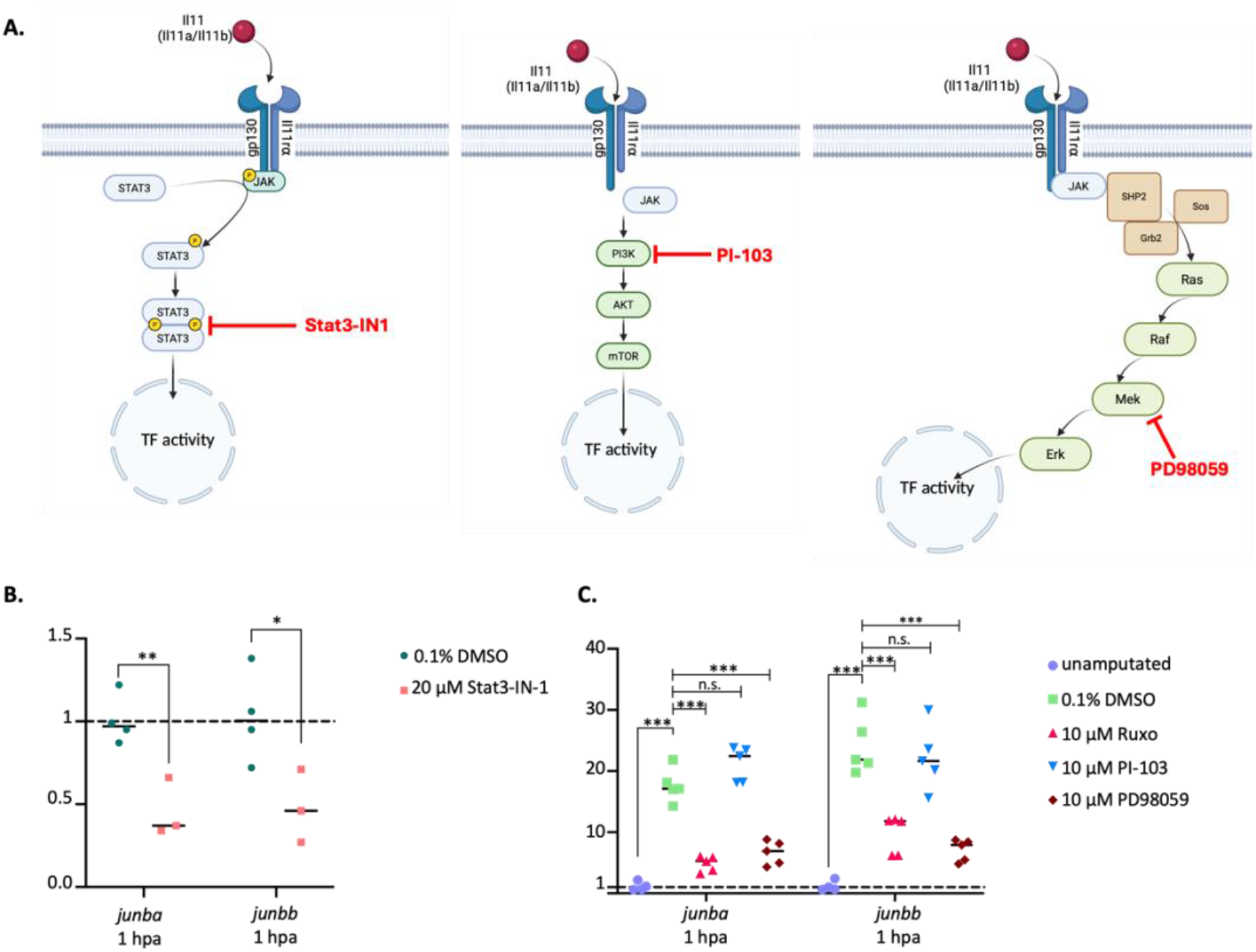
The expression of *junba* and *junbb* depends on the Stat3 pathway of Interleukin 11 signalling. **A.** Schematic depiction of the different paths of IL-11 signalling and inhibitors than can be used to specifically inhibit the pathways. The small molecule compound inhibitor prevents the Stat3 path of IL-11 signalling. The pyridinylfuranopyrimidine compound PI-103 inhibits the PI3K-mTOR path of IL-11 signalling. The small molecule compound inhibitor PD98059 prevents activation of the MEK/ERK dependent expression of IL-11 target genes. **B.** Analysis of *junba* and *junbb* expression in larvae treated with the the Stat3 Inhibitor Stat3-IN1 compared to the vehicle control (n= 4 tissue, each sample from 20 pooled larvae), at 1hpa. Significance is indicated as *p<0.05 and **p<0.01. **C.** Analysis of *junba* and *junbb* expression from larvae treated with Ruxolitinib, PI-103 & PD98059 compared to the vehicle control at 1hpa. (n = 4 tissue, each sample from 20 pooled larvae). Significance is indicated as ***p<0.001.

**Supplementary Figure 3:**
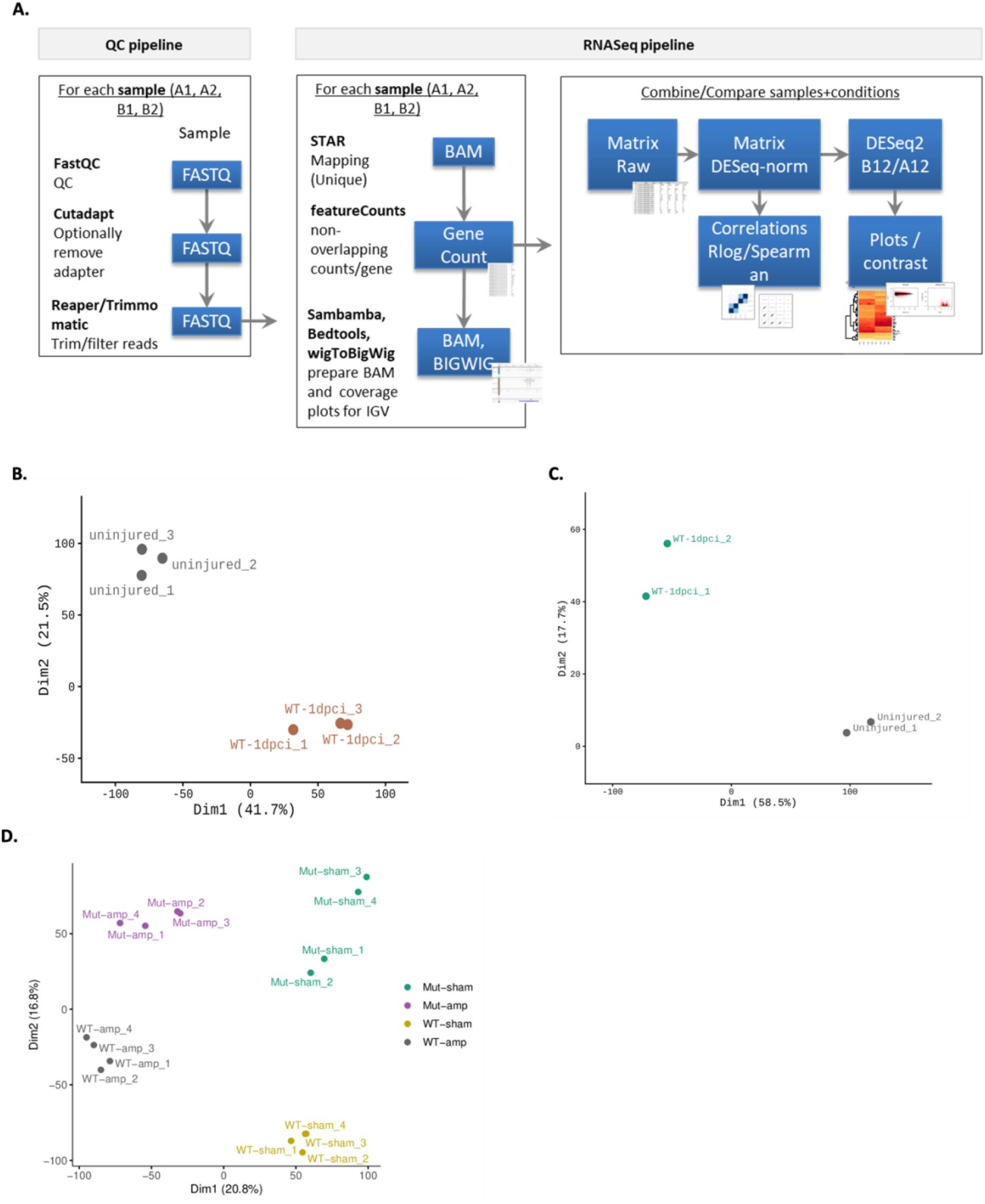
Pipeline of bulk RNAseq data processing. (B-D) Principal Component Analysis (PCA). Downstream analyses were based on the normalized gene count matrix. A global clustering heatmap of samples was created based on the euclidean distance of regularized log transformed gene counts. Dimension reduction analyses (PCA) were performed on regularized log transformed counts using the R packages FactoMineR (Le et al., FactoMineR: A Package for Multivariate Analysis). (B) PCA of mouse heart samples namely, uninjured and 24h post injury (WT-1dpci) (C) PCA of zebrafish heart samples namely, uninjured and WT-1dpci (24h post injury (D) PCA of WT and *junb* ^-/-^ dm (Mut) larvae. Here uninjured is denoted by sham and injured by (amp).

## Notes

### Competing Interest Statement

The authors have declared no competing interest.

